# Adaptive Digital Tissue Deconvolution

**DOI:** 10.1101/2023.02.08.527583

**Authors:** Franziska Görtler, Malte Mensching-Buhr, Ørjan Skaar, Stefan Schrod, Thomas Sterr, Andreas Schäfer, Tim Beißbarth, Anagha Joshi, Helena U. Zacharias, Sushma Nagaraja Grellscheid, Michael Altenbuchinger

## Abstract

**Motivation:** The inference of cellular compositions from bulk and spatial transcriptomics data increasingly complements data analyses. Multiple computational approaches were suggested and recently, machine learning techniques were developed to systematically improve estimates. Such approaches allow to infer additional, less abundant cell types. However, they rely on training data which do not capture the full biological diversity encountered in transcriptomics analyses; data can contain cellular contributions not seen in the training data and as such, analyses can be biased or blurred. Thus, computational approaches have to deal with unknown, hidden contributions. Moreover, most methods are based on cellular archetypes which serve as a reference; e.g., a generic T-cell profile is used to infer the proportion of T-cells. It is well known that cells adapt their molecular phenotype to the environment and that pre-specified cell archetypes can distort the inference of cellular compositions.

**Results:** We propose Adaptive Digital Tissue Deconvolution (ADTD) to estimate cellular proportions of pre-selected cell types together with possibly unknown and hidden background contributions. Moreover, ADTD adapts prototypic reference profiles to the molecular environment of the cells, which further resolves cell-type specific gene regulation from bulk transcriptomics data. We verify this in simulation studies and demonstrate that ADTD improves existing approaches in estimating cellular compositions. In an application to bulk transcriptomics data from breast cancer patients, we demonstrate that ADTD provides insights into cell-type specific molecular differences between breast cancer subtypes.

**Availability and implementation:** A python implementation of ADTD and a tutorial are available at Gitlab and zenodo (doi:10.5281/zenodo.7548362).

**Supplementary information:** Supplementary material is available at *Bioinformatics* online.

## Introduction

Bulk transcriptomics profiles are a complex linear combination of diverse molecular contributions of multiple individual cells. Many analyses, such as the screening for differentially expressed genes, can be confounded by the underlying cellular compositions. Consequently, if compositions are unknown, we cannot resolve the source of differential gene expression. Multiple approaches were suggested to estimate cellular proportions from bulk transcriptomics data, ranging from traditional, regression-based approaches to machine-learning techniques, including generalized regression (Chen *et al*., 2018; Altboum *et al*., 2014; Du *et al*., 2019) as well as deep learning approaches (Menden *et al*., 2020; Lin *et al*., 2022). In recent years, the advent of spatial transcriptomics has generated an additional need for reliable cell-type deconvolution (Dong and Yuan, 2021; Ma and Zhou, 2022).

Single-cell RNA sequencing data can substantially improve estimates of cellular proportions via improved reference matrices as well as collections thereof, and via optimized gene selection or weighting, as demonstrated by work of us and others (Tsoucas *et al*., 2019; Wang *et al*., 2019; Görtler *et al*., 2020; Jew *et al*., 2020; Dong *et al*., 2021). The latter can be facilitated via artificial mixtures of known cellular composition generated from single-cell data. For instance, MuSiC (Wang *et al*., 2019) and Digital Tissue Deconvolution (DTD) (Görtler *et al*., 2020) learn a gene weighting, and Bisque (Jew *et al*., 2020) learns gene-specific bulk expression transformations.

A recent comprehensive benchmarking of different methods for cell-type deconvolution resolved the marker gene selection as a major determinant of model performance and it was pointed out that if cell types are missing in the reference profiles results become substantially worse (Avila Cobos *et al*., 2020). The latter statement is in line with the observation that MuSiC, Non-Negative Least Squares (NNLS), and CIBERSORT (Chen *et al*., 2018) did not produce accurate results in a validation study using artificial bulk data of six pancreatic cell types if one cell type was removed from the reference single-cell expression dataset (Wang *et al*., 2019). Consequently, all cell types which are contained in the bulk mixtures should ideally also be included in the reference matrix (Avila Cobos *et al*., 2020). This, however, represents a major issue in practical applications, since bulk profiles likely contain signals of cell types not originally considered for model development.

Another, related issue is the origin of reference profiles. It is well known that the cell’s molecular phenotype depends on its environment. This also affects the results of digital tissue dissection, as shown by (Schelker *et al*., 2017; Racle *et al*., 2017), where it was pointed out that the references should be selected from niches similar to the investigated bulks.

Although many different approaches were meanwhile suggested for cell-type deconvolution, they rarely account for hidden cell type contributions and cellular environmental effects. Here, we systematically address both issues within a single approach named Adaptive Digital Tissue Deconvolution (ADTD). First, ADTD builds on our previous work on DTD and uses gene weights for optimized deconvolution. Second, it augments the deconvolution by estimates of a background profile together with corresponding cellular proportions across a set of investigated bulks. Third, ADTD adapts its reference profiles to the investigated bulks. Intriguingly, the latter also resolves cell-type specific regulation from bulk transcriptomics data. We verified ADTD in simulations based on single-cell RNA sequencing data, illustrating that ADTD out-competes state-of-the-art baselines. This particularly holds true if models are transferred to another domain, as shown for the transfer from healthy breast tissue to breast cancer. We finally illustrate how ADTD complements routine data analysis in an application to bulk transcriptomics data from The Cancer Genome Atlas (TCGA; https://www.cancer.gov/tcg) revealing patterns of cell-type specific gene regulation in breast cancer.

## Methods

Adaptive digital tissue deconvolution (ADTD) builds on loss-function learning for DTD introduced previously by Görtler *et al*., 2020 and extends it by both accounting for unknown background contributions and adaptive reference profiles. We will first recapitulate DTD and will then introduce ADTD.

Loss-function learning for digital tissue deconvolution Bulk gene expression profiles can be modeled as a linear combination of cell-type-specific reference profiles,

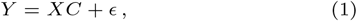

Where 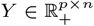 is a matrix with *n* bulk gene expression profiles and 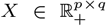 a matrix with *q* cell-type specific reference profiles in their columns, both containing expression values of *p* genes in their rows. The columns of 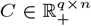 contain cellular weights corresponding to the columns (bulks) of *Y* . The values *C*_*ki*_ can be interpreted as semi-quantitative measurements of the number of cells of type *k* in bulk profile *i*. The residuals between the observed bulks *Y* and the fitted bulks *Ŷ* = *XĈ* are given by *ϵ* ∈ ℝ^*p×n*^.

One possible way to estimate *C* is by

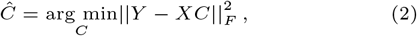

where ||.||_*F*_ is the Frobenius norm and *Ĉ* the estimate of *C* for given *X* and *Y* . Eq. (2) can be solved for each column (bulk) in *Y*, individually. In (Görtler *et al*., 2020; Schön *et al*., 2020), Eq. (2) was augmented by gene weights *g* = (*g*_1_, *g*_2_, …, *g*_*p*_)^*T*^, yielding

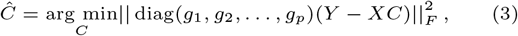

where the weights *g*_*i*_ can be chosen to improve estimates of *C*. They can be either specified by prior knowledge or can be learned from cellular mixtures with known cellular contributions, such as annotated bulk profiles or artificial mixtures generated from single-cell RNA sequencing data (Görtler *et al*., 2020; Schön *et al*., 2020). To optimize the gene weights, we defined the outer loss function

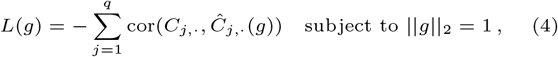

where *C*_*j,·*_ is the *j*th row (cell type) of *C* and *Ĉ*_*j,·*_ its corresponding estimate. Thus, minimizing loss function Eq. (4) with respect to the *g*_*i*_s, maximizes the Pearson’s correlation between the ground truths *C*_*j,·*_ and its estimates *Ĉ*_*j,·*_ for all cell types of interest *j*, simultaneously. In contrast to the original implementation of loss-function learning (Görtler *et al*., 2020), we now additionally restrict the optimization problem to non-negative (physical) solutions *Ĉ* ⪰ 0_*q,n*_. The corresponding algorithm uses the PyTorch machine learning library (Paszke *et al*., 2019) with stochastic gradient descent for optimization.

### Estimates of background contributions

One major drawback of reference-based cell-type deconvolution methods is that they do not necessarily capture all cellular contributions present in complex biospecimens. For instance, let *X* consist of a set of immune cell reference profiles and let *Y* be the bulk profile of a tumor specimen. Then, although immune cell contributions can be estimated, their values might be biased or blurred by the contributions from cancer cells to *Y* . In seminal work by Racle *et al*., 2017, this issue was addressed by a method called EPIC (Estimating the Proportion of Immune and Cancer cells), as outlined in the following. Let *Y*_*·,i*_ be an individual bulk profile corresponding to the *i*th column of *Y* . Then, we have

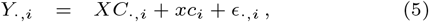

where we made the contribution from a potential background explicit by adding the term *xc*_*i*_. Here, *x* ∈ ℝ ^*p×*1^ represents the “hidden” background profile and *c*_*i*_ its corresponding scalar weight. We further assume that the bulk profiles, the reference profiles, and the background profile are normalized by

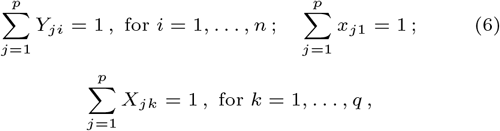

which gives 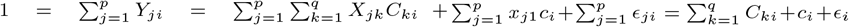 with 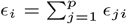. This implies 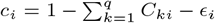 and motivates that *c*_*i*_ can be directly estimated as

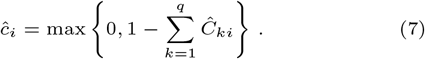

Thus, EPIC intrinsically assumes that *C* can be estimated independently from the background profile *x* and that all *ϵ*_*i*_ are small.

### Adaptive Digital Tissue Deconvolution

#### Background estimation

ADTD relaxes EPIC’s assumptions and directly takes into account that the inference of *C* also depends on the background profile *x*. To achieve this, we replace *c*_*i*_ in the residuals 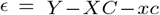 of Eq. (5) by 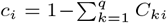 (assuming that the *ϵ*_*i*_s are small), and minimize their squared sum with respect to *C* and *x*. The corresponding optimization problem taking into account the gene weights determined via loss-function learning and non-negativity constraints becomes

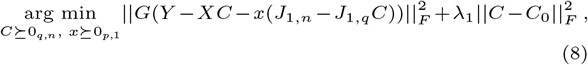

subject to *J*_1,*q*_*C* ⪯ *J*_1,*n*_ and 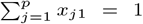, where *G* = diag(*g*_1_, *g*_2_, …, *g*_*p*_), *C*_0_ is the estimate of *C* using an ordinary NNLS estimate weighted by *γ*_*j*_ = *g*^2^, and *J*_*a,b*_ denotes a matrix of ones of dimension *a* × *b*. The regularization parameter *λ*_1_ calibrates between two solutions. For *λ*_1_ → ∞, the naive solution is retrieved, where the hidden cell proportions are given by Eq. (7). For *λ*_1_ → 0, *x* and *C* are estimated simultaneously. With growing *λ*_1_ values, the solutions are more strongly biased toward the naive solution *Ĉ* = *C*_0_. Note that the naive solution can be determined for each cellular mixture individually, while the latter solution requires a sufficient amount of bulk samples, however, providing the advantage that a consensus background is learned across a set of bulk specimens. We solved the optimization problem Eq. (8) by iteratively optimizing with respect to *C* and *x* using quadratic programming, until convergence is reached.

#### Adaptive reference profiles

Reference profiles can be generated using external data from purified cellular mixtures or from single-cell data. However, cells adapt their molecular phenotype to their environment and as a consequence, global and static reference profiles might be inappropriate for most applications. ADTD dynamically adapts reference profiles to the specific application in order to improve inferred cellular contributions and to reveal potential regulatory effects on cell populations. This is achieved by replacing the reference matrix *X* by Δ ◦ *X*, where Δ is a *p* × *q* matrix of rescaling factors which are applied component-wise to *X* (◦ is the Hadamard product). The full ADTD loss function is defined as

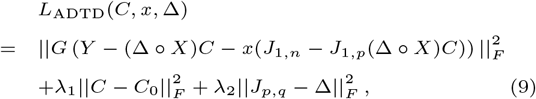

which is optimized with respect to *C, x*, and Δ, subject to the constraints

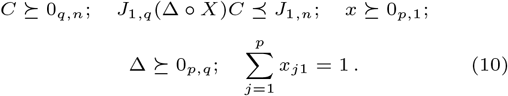

In Eq. (9), we replaced 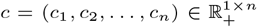 by *J*_1,*n*_ − *J*_1,*p*_(Δ ◦ *X*)*C* as derived in Supplementary Section 1.1. The regularization term 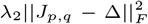 controls overfitting with respect to the parameters Δ_*jk*_. For *λ*_2_ → ∞, we get Δ = *J*_*p,q*_. Thus, the limit *λ*_2_ → ∞ corresponds to ADTD without cellular adaptation. Further *C*_0_ is determined via DTD with non-negativity constraint. Thus, the DTD limit is retrieved for *λ*_1_ → ∞.

#### Implementation

The ADTD optimization problem was solved by iteratively optimizing the loss *L*_ADTD_(*C, x*, Δ) with respect to *C, x*, and Δ until convergence is reached. The minimization with respect to *C* and *x* was addressed by quadratic programming (see Suppl. Sections 1.2 and 1.3). The optimization with respect to Δ was performed iteratively for the rows Δ_*j,·*_ (Suppl. Section 1.4). The full algorithm is summarized below (Algorithm 1).

##### Algorithm 1 Adaptive Digital Tissue Deconvolution

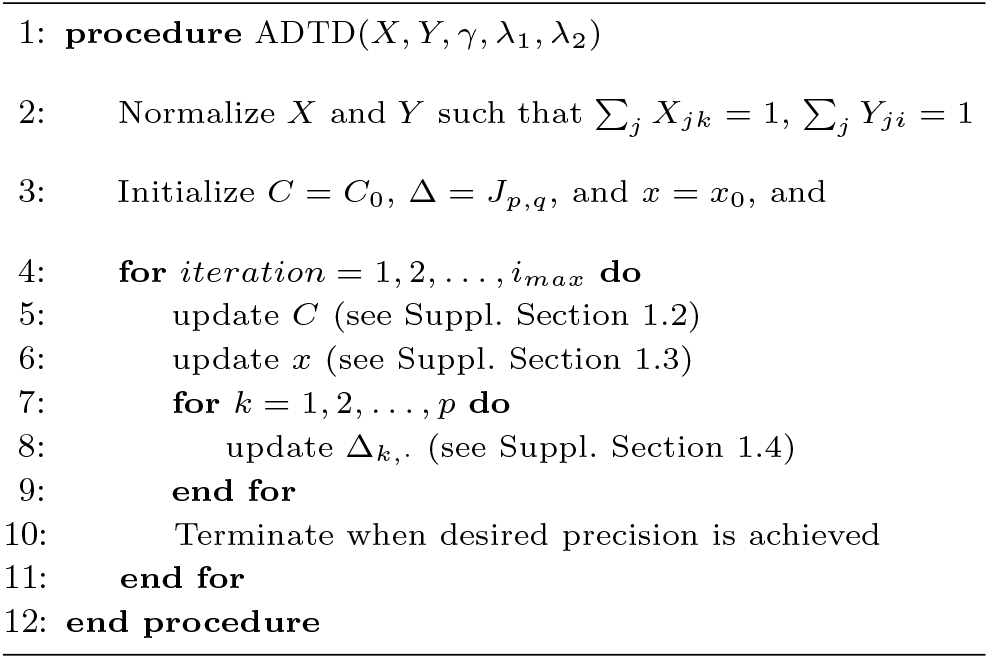

### Data processing, simulation studies, and analyses

#### Artificial bulks from single-cell data of healthy breast tissue – training data

We generated artificial mixtures of known cellular compositions for model training using single-cell data from healthy breast tissue (GSE164898) (Bhat-Nakshatri *et al*., 2021), representing women of different race, age, parity, menstrual phase, Tyrer-Cuzick score, and Body Mass Index (BMI). We denote these data as “training data” throughout the article. Cell types were labeled using Seurat (Hao *et al*., 2021), following the workflow of (Wu *et al*., 2021) to remain consistent with the test data described below. We averaged across all respective single-cell profiles of individual patients to derive archetypic reference profiles of B-cells, endothelial cells, myeloid cells, epithelial cells, perivascular-like cells (PVL), and T-cells, yielding a reference matrix for each patient. These were subsequently averaged to yield a global reference matrix *X*. Cells which were not labeled as one of these six cell types were removed from the training set. We further reduced the gene space to in total 1000, as follows. We first included the top 30 genes of every Seurat main cluster. Of the main clusters, we further investigated those containing predominantly endothelial cells and mesenchymal cells. These were re-clustered to yield 18 sub-clusters and of each of those, we further included the top 10 genes. This yielded in total 524 unique genes (genes representing multiple cluster were included only once). We further ranked all genes according to their variance calculated with respect to *X* and selected the 476 most variable genes to yield in total 1000 selected genes. We generated individual mixtures (*n* = 5000) by randomly drawing 100 single-cell profiles each and by averaging their expression.

#### Artificial bulks from single-cell data of healthy breast tissue – validation data

For a first validation of ADTD, we generated validation mixtures from the same single-cell profiles as previously, but now also using the hold-out single-cell profiles; the mixtures now include hidden background contributions from cells not assigned to one of the six major cell types (∼ 15% of all cells). To also study ADTD’s capability to reconstruct cell-type specific regulation from bulk transcriptomics data, we further artificially modified gene-expression levels in a cell-type specific way. For this, we modified the expression of gene *j* in mixture *i* by (1) drawing a modification factor *a*_*i*_ from the set {−1, −0.5, 1, 2}, followed by (2) adding *a*_*i*_*X*_*kj*_ *C*_*ki*_ to the original expression *Y*_*ij*_ (values *Y*_*ij*_ *<* 0 were set to zero), where *k* corresponds to the cell profile. In each simulation run, we modified 60 genes; 30 genes randomly drawn from the 20 most specific genes for each cell type (in total 120 genes, selected via max_*k*_ {*X*_*jk*_*/* mean(*X*_*j,·*_)}) and 30 genes randomly drawn from the 120 most unspecific genes (selected according to minimal values of var(*X*_*j,·*_)*/* mean(*X*_*j,·*_)). Thus, we manipulated both genes which are particularly relevant for a given cell type and genes which are rather unspecific for cell types. We refer to the so generated data as “validation data” throughout the article.

#### Artificial bulks from single-cell data of breast cancer tissue – test data

Artificial test mixtures of known cellular compositions were generated analogously to the training data, but now using single-cell data of breast cancer specimens (Wu *et al*., 2021). Thus, these data specifically allow us to study the domain transfer to completely unseen data, generated within a different experiment, and capturing a substantially different tissue as a consequence of cancer cell admixtures. In total, single-cell data of 26 different tumors were available with corresponding cell-type labels. We extracted B-cells, endothelial cells, myeloid cells, epithelial cells, PVL cells, and T-cells, as described above, and additionally cancer epithelial cells. These profiles were used to generate *n* = 5000 artificial bulk mixtures by randomly drawing 100 single-cell profiles each. In this process, we further added cell-type specific gene regulation, as previously done for the healthy validation data.

#### TCGA breast cancer analysis

We retrieved the BRCA bulk transcriptomics data generated by the TCGA Research Network (The Cancer Genome Atlas, https://www.cancer.gov/tcga), containing, in total, 1083 samples: 190 triple-negative breast cancers (TNBC; ER−, PR−, HER2−), 82 human epidermal growth factor receptor 2-positive breast cancer (HER2+), 562 luminal A (LumA; ER+, PR+, HER2−, and low levels of Ki-67), 209 luminal B (LumB; ER+, PR+*/*− and HER2+), and 40 breast cancer free tissue samples (normal). To compensate systematic differences between the measurement technologies, we further normalized the TCGA data gene-wise to the single-cell training data. Gene-wise re-scaling factors were calculated by dividing the respective mean expression in the single cell pseudo bulks of the training set by those derived for the TCGA controls.

#### Competing methods

We compared ADTD to several competing methods. First, we compared it to two different implementations of EPIC (Racle *et al*., 2017), one using our own reference matrix *X* generated from the training data (EPIC_1_) and the originally suggested version using the EPIC built-in reference matrix (EPIC_2_). EPIC_2_ returns only estimates for B-cells, endothelial cells, macrophages, PVLs, and T-cells. Therefore, we associated estimated macrophage contributions of EPIC_2_ with the ground truth of myeloid lineage cells. Second, we compared it to CIBERSORTx (Newman *et al*., 2019), where we used our own reference matrix, since the built-in matrix did not resemble the herein investigated cell types sufficiently. Third, we compared it to Scaden, which is a deep-learning based approach to estimate cellular proportions. Thus, similar as we trained the DTD gene weighting, we learned Scaden to disentangle the training data. For the latter, healthy single-cell profiles of our training data were provided for model development.

## Results

### Model training and validation

#### Model development and competing methods

It was repeatedly shown that gene selection strongly affects the performance of bulk deconvolution methods (Görtler *et al*., 2020; Racle *et al*., 2017), and the importance of a stringent marker gene selection was thoroughly demonstrated (Avila Cobos *et al*., 2020). ADTD involves a two step procedure. In a first step, it uses DTD (Görtler *et al*., 2020) on the training data to optimize cell-type deconvolution via gene weights *g*_*i*_ in Eq. (3). Thus, DTD does not select marker genes but weighs them. Here, DTD was trained on the training data to disentangle B-cells, endothelial cells, myeloid cells, normal epithelial cells, PVLs, and T-cells. Thus, to this point, the training process did not see any cellular background and all cells in the mixtures are also represented by the reference matrix *X*. Suppl. Table S1 summarizes the training performance for ADTD and the competing methods, where we evaluated the models by drawing *n* = 1000 samples. The results show that both ADTD and Scaden perform substantially better than the competitors. However, both were optimized to fit the data while the others were not.

##### ADTD hyper-parameters have little effect on training performance

ADTD requires specification of the hyper-parameters *λ*_1_ and *λ*_2_. While those do not affect the DTD prior learning, they affect predictions. We evaluated a comprehensive parameter grid consisting of all combinations of *λ*_1_ ∈ {0, 10^−5^, 10^−4^, …, 1, 10, ∞} with *λ*_2_ ∈ {10^−9^, 10^−8^, …, 10, ∞}, showing that ADTD is little sensitive to the explicit choice of the regularization terms, including the solutions with *C* = *C*_0_ for *λ*_1_ → ∞ and Δ = *J*_*p,q*_ for *λ*_2_ → ∞, which also performed remarkably well (Suppl. Fig. S1). The only exception is when both regularization parameters are extremely small *<* 10^−7^, simultaneously, or when *λ*_1_ = 0. The ADTD results in Suppl. Table S1 correspond to *λ*_1_ = 10^−1^ and *λ*_2_ = 10^−8^. This explicit choice was motivated using the validation data and subsequently verified on the independent breast cancer test data, as outlined in the following.

#### Validation performance

In a next step, we verified ADTD’s and the competitors’ performance on the artificial validation data. Those were generated from the same measurement batch (healthy tissue) but additionally included contributions from single-cell profiles which could not be assigned to any of the major cell clusters and as such represent a cellular background (see methods). Deconvolution algorithms have to deal with such unknown cellular contributions and among the investigated methods, only ADTD and EPIC are able to estimate background contributions, while CIBERSORTx and Scaden are not. Nevertheless, the latter two were included as additional state-of-the-art baselines for comparison.

Table 1 gives the performance of ADTD and the competitors in estimating the ground truth cellular compositions for each of the cell types included in the reference matrix for *n* = 1000 mixtures. We observe that ADTD translates best to these first validation data with minimal performance losses compared to the training data (mean Pearson’s correlation over all cell types; 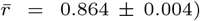. ADTD is followed by Scaden 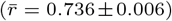, and both perform substantially better than 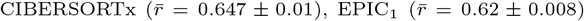 and 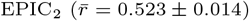, respectively.

**Table 1.** Performance of ADTD, EPIC, CIBERSORTx and Scaden on validation data (healthy tissue). Observed Pearson’s correlations obtained by comparing estimated with true cellular proportions for artificial mixtures generated from single-cell data of healthy breast tissue (see Methods). The error bars correspond to ±1 standard deviation obtained over 10 simulation runs. Correlations *>* 0.8 are highlighted. Abbreviations: Endo. = endothelial cells; Myel. = myeloid cells; Epith. = epithelial cells; PVL = perivascular-like cells.

|  | ADTD | EPIC<sub>1</sub> | EPIC<sub>2</sub> | CIBERSORTx | Scaden |
| --- | --- | --- | --- | --- | --- |
| B-cells | 0.744 ± 0.015 | 0.094 ± 0.034 | 0.419 ± 0.028 | 0.126 ± 0.037 | 0.551 ± 0.020 |
| Endo. | **0.908 ± 0.003** | **0.839 ± 0.008** | 0.712 ± 0.017 | **0.824 ± 0.009** | **0.830 ± 0.009** |
| Myel. | **0.929 ± 0.004** | **0.825 ± 0.006** | 0.678 ± 0.018 | **0.835 ± 0.018** | **0.801 ± 0.013** |
| Epith. | **0.906 ± 0.007** | 0.647 ± 0.010 | - | 0.760 ± 0.019 | 0.781 ± 0.019 |
| PVL | 0.773 ± 0.016 | 0.530 ± 0.026 | - | 0.569 ± 0.013 | 0.693 ± 0.020 |
| T-cells | **0.925 ± 0.003** | 0.782 ± 0.011 | 0.284 ± 0.032 | 0.768 ± 0.012 | 0.759 ± 0.015 |
| mean | **0.864 ± 0.004** | 0.620 ± 0.008 | 0.523 ± 0.014 | 0.647 ± 0.010 | 0.736 ± 0.006 |

We further evaluated ADTD and EPIC_1,2_ with respect to their performance in inferring the hidden cellular background (Table 2). Respective cellular proportions were estimated with a Pearson’s correlation of *r* = 0.840 ± 0.010 for ADTD and *r* = 0.262±0.014 for EPIC_1_. Here, EPIC_2_ was not evaluated since it captures a different set of cell types via *X*, which also affects the reconstructed backgrounds. As such, a fair comparison would not be possible. Additionally, ADTD provides estimates of the hidden-cell profile *x*, which could be also reliably reconstructed (*r* = 0.754 ± 0.007).

**Table 2.** Performance of ADTD and EPIC_1_ in estimating hidden contributions on the healthy validation data. Performance in terms of Pearson’s correlation for the estimated hidden proportions (hidden cell type) and the hidden profile.

|  | ADTD | EPIC<sub>1</sub> |
| --- | --- | --- |
| hidden cell type | **0.840 ± 0.010** | 0.262 ± 0.014 |
| hidden profile | 0.754 ± 0.007 | - |

##### ADTD hyper-parameter dependency

As for the training data, we systematically evaluated ADTD’s performance for different choices of *λ*_1_ and *λ*_2_ with results shown in Figure S2a and b, providing the average performance for estimating cellular proportions of the included cell types (Fig. S2a) and the hidden cells (Fig. S2b). We confirmed a negligible dependence of the ADTD’s prediction performance with respect to *λ*_1_ and *λ*_2_. As previously on the training data, the only exception is the case when both *λ*_1_ and *λ*_2_ are very small, simultaneously.

##### Reverse engineering of cell-type specific gene regulation and hyper-parameter dependency

in contrast to the competing methods, ADTD adapts the reference matrix via re-scaling factors encoded by Δ. Our artificial validation mixtures were generated to contain such effects, resembling potential cell-type specific gene-regulatory contributions to gene expression. We verified that ADTD is able to resolve cell-type specific gene regulation by evaluating areas under the Receiver Operating Characteristic (AUC) curves. As previously, we evaluated ADTD for different choices of *λ*_1_ and *λ*_2_ (Suppl. Fig. S2c). In contrast to the previous prediction tasks, an appropriate choice of the hyper-parameters turns out to be much more important in this context. We observed that for large values of *λ*_2_ the AUCs approach 0.5, corresponding to random predictions. This might be expected, since Δ = *J*_*p,q*_ for *λ*_2_ → ∞. For smaller values of *λ*_2_, ADTD increasingly shows its capability to resolve cell-type specific gene expression, yielding AUCs up to ∼ 0.9. Importantly, performance is little dependent on the explicit choice of *λ*_2_, providing reasonable performance in the full range from 10^−9^ (AUC∼ .9) to 10^−3^ (AUC∼ .8) given that *λ*_1_ *>* 0.3. Again, if both are very small, performance is compromised, indicating potential issues with over-fitting. This is also supported by Fig. S3 and S4, where performance with respect to *λ*_2_ is studied in detail for the two explicit choices *λ*_1_ = 10^−1^ and *λ*_1_ = 10^−3^.

##### Hyper-parameter selection

given the outlined performance evaluations, we decided to fix *λ*_1_ = 10^−1^ and *λ*_2_ = 10^−8^, since for this choice validation performance is good with respect to all investigated measures. The performance for alternative choices is given in S2a to c (validation data) and S2d to f (test data, and see below).

#### Domain transfer to breast cancer

Next, we systematically studied the domain transfer to breast cancer tissue. For this purpose, we generated artificial cellular mixtures using single-cell data from breast cancer specimens (Wu *et al*., 2021). Thus, the test mixtures are generated from a different single-cell experiment and address a different tissue context, namely breast tissue infiltrated by cancer cells. Thus, the artificial mixtures contain a substantial proportion of cells not seen in the training data.

##### Performance comparison to competing methods

Table 3 summarizes the results for ADTD and the competing methods. We observe a substantially better performance of ADTD compared to the competitors for each individual cell type, with the only exception of normal epithelial cells. The latter were slightly better estimated by Scaden (*r* = 0.349 ± 0.022 for ADTD vs. *r* = 0.378 ± 0.03 for Scaden). ADTD’s average performance was 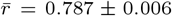, followed by Scaden 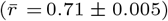, EPIC_2_ 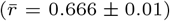, EPIC_1 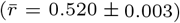_, and CIBERSORTx 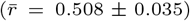. Considering estimates of hidden cellular contributions (see Table 4), ADTD performs clearly better than EPIC_2_ (*r* = 0.449 ± 0.031 for ADTD vs. *r* = 0.24 ± 0.017 for EPIC_2_). For the outlined reasons, a comparison to other approaches was not possible here. One exemplary scatter plot contrasting ADTD predictions with the corresponding ground truth is shown in Suppl. Fig. S5. The comparatively low performance for estimating epithelial and cancer epithelial contributions might be explained by the fact that both belong to a continuum of cells, consisting of cancer cells, epithelial cells, but also a number of boundary cases which resemble molecular characteristics of both. To determine the ground truth, we used the labelling provided by (Wu *et al*., 2021) based on a scoring system to annotate cancerous cells (the genomic instability score).

**Table 3.** Performance of ADTD, EPIC, CIBERSORTx and Scaden on the breast cancer test data. Observed Pearson’s correlations obtained by comparing the estimated cellular proportions of the cell types captured in the reference matrix *X* with the ground truth for artificial cellular mixtures of the breast cancer test data set, where solely the cancer epithelial cells were hidden in the mixtures and not represented as reference profiles (see also Table 4). The error bars correspond to ±1 standard deviation obtained over 10 simulation runs and correlations *>* 0.8 are highlighted. Abbreviations: Myel. = myeloid cells; Epith. = epithelial cells; PVL = perivascular-like cells.

|  | ADTD | EPIC<sub>1</sub> | EPIC<sub>2</sub> | CIBERSORTx | Scaden |
| --- | --- | --- | --- | --- | --- |
| B-cells | **0.824 ± 0.012** | 0.035 ± 0.022 | 0.717 ± 0.013 | 0.315 ± 0.089 | 0.657 ± 0.018 |
| Endo. | **0.832 ± 0.008** | 0.631 ± 0.011 | 0.783 ± 0.010 | 0.633 ± 0.060 | **0.859 ± 0.010** |
| Myel. | **0.928 ± 0.003** | **0.867 ± 0.005** | 0.748 ± 0.012 | 0.798 ± 0.044 | 0.794 ± 0.009 |
| Epith. | 0.349 ± 0.022 | 0.281 ± 0.023 | - | 0.201 ± 0.092 | 0.378 ± 0.030 |
| PVL | **0.907 ± 0.005** | 0.573 ± 0.014 | - | 0.670 ± 0.062 | 0.838 ± 0.007 |
| T-cells | **0.882 ± 0.007** | 0.732 ± 0.018 | 0.419 ± 0.035 | 0.427 ± 0.096 | 0.732 ± 0.015 |
| mean | 0.787 ± 0.006 | 0.520 ± 0.003 | 0.666 ± 0.010 | 0.508 ± 0.035 | 0.710 ± 0.005 |

**Table 4.** Performance of ADTD and EPIC_1_ in estimating hidden contributions on the breast cancer test data. Performance in terms of Pearson’s correlation for the estimated hidden proportions (cancer Epith.) and the hidden profile.

|  | ADTD | EPIC<sub>1</sub> |
| --- | --- | --- |
| cancer Epith. | 0.449 ± 0.031 | 0.24 ± 0.017 |
| hidden profile | 0.712 ± 0.004 | - |

##### Sample-size dependency

ADTD solves an optimization problem which simultaneously estimates the hidden reference profile *x* and the cellular proportions. Thus, an accurate estimate of *x* requires accurate estimates of *C* and vice versa. The hidden background profile *x* can be considered as an average background which is observed among different mixtures. Thus, to obtain reliable estimates for *x* and *c*, one might need to consider multiple mixtures. To study the impact of sample size *n* on ADTD test performance, we repeated the performance evaluation for different numbers of samples with results shown in Figure 1, where the observed Pearson’s correlation is plotted versus sample size *n* for the cell types contained in the reference matrix (Figs. 1a to f), and for the estimated hidden proportions (Fig. 1g). Additionally, Fig. 1h shows the correlations comparing the estimated background profile *x* with the ground truth. Strikingly, ADTD achieves already remarkable stable performance for sample sizes as low as *n* = 50.

**Fig. 1.**
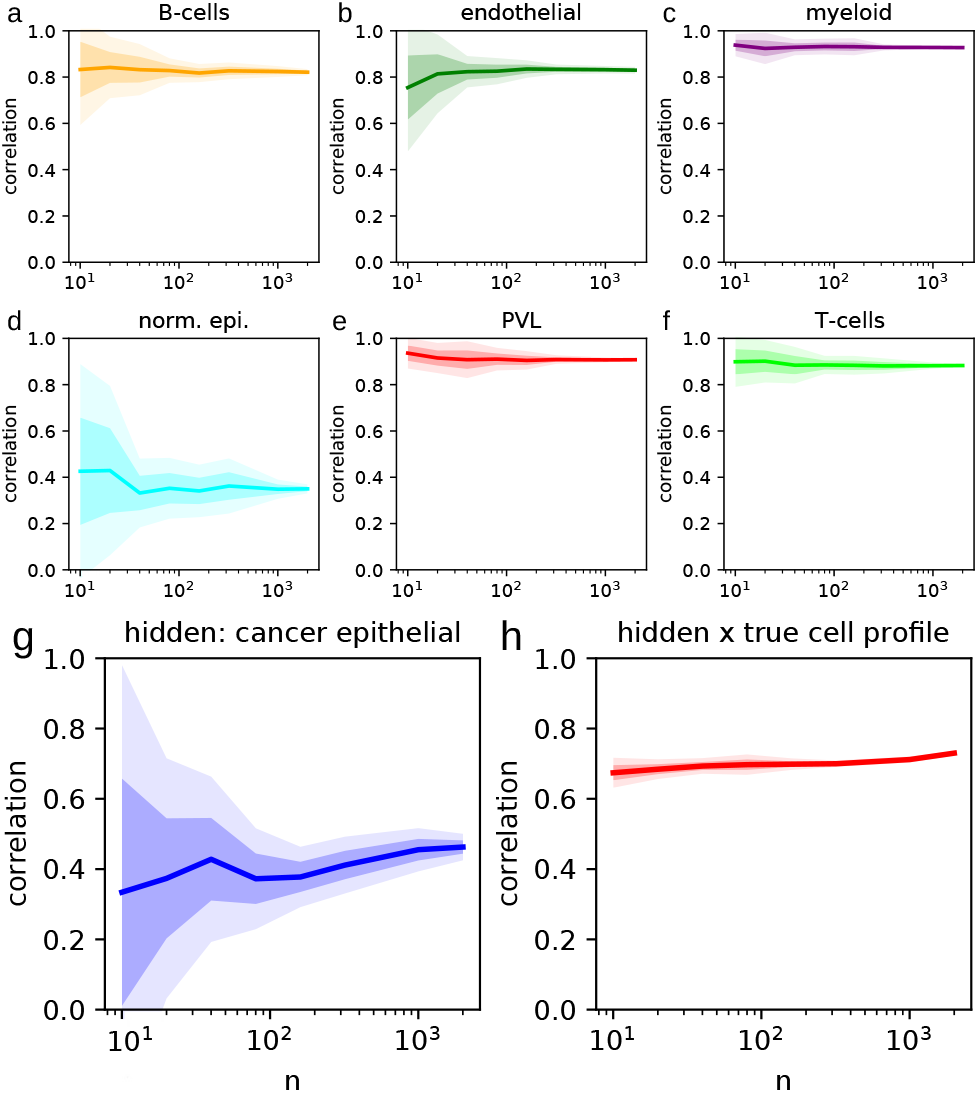
Simulation illustrating sample-size stability of ADTD on the test data. Performance versus sample size *n* for ADTD in terms of Pearson’s correlation for comparing estimates with their ground truth. Figures a to f show the performance for cell types captured in the reference matrix *X*, corresponding to B-cells, endothelial cells, myeloid cells, normal epithelial, PVLs, and T-cells, respectively. Figure g shows the corresponding *n* dependency for the hidden proportion, corresponding to cancer epithelial cells. Figure h shows the Pearson’s correlation for comparing the estimated background profile *x* of cancer epithelial cells with the ground truth. The dark shaded bands correspond to *±*1 standard deviation and the light shaded bands to *±*2 standard deviations calculated across 10 simulation runs.

##### Reconstruction of cellular regulation

We further assessed ADTD’s performance to reverse engineer cell-type specific gene regulation in our breast cancer test mixtures. Figure 2a provides respective performance evaluation in terms of AUCs for different values of *λ*_2_ for a fixed value of *λ*_1_ = 10^−1^, containing also the finally selected model (*λ*_1_ = 10^−1^, *λ*_2_ = 10^−8^). ADTD remains highly predictive for the whole range of *λ*_2_ values between 10^−9^ to 10^−4^, with best results observed for 10^−9^, closely followed by 10^−8^. Suppl. Fig. S6 shows a similar behaviour for *λ*_1_ = 10^−3^. We further tested how performance depends on sample size with results shown in Fig. S7, where we observed remarkably stable and still reasonably high performance for mixtures with sample sizes of only *n* = 125. This was further substantiated for one alternative model (*λ*_1_ = 10^−3^, *λ*_2_ = 10^−6^) in Suppl. Fig. S8.

**Fig. 2.**
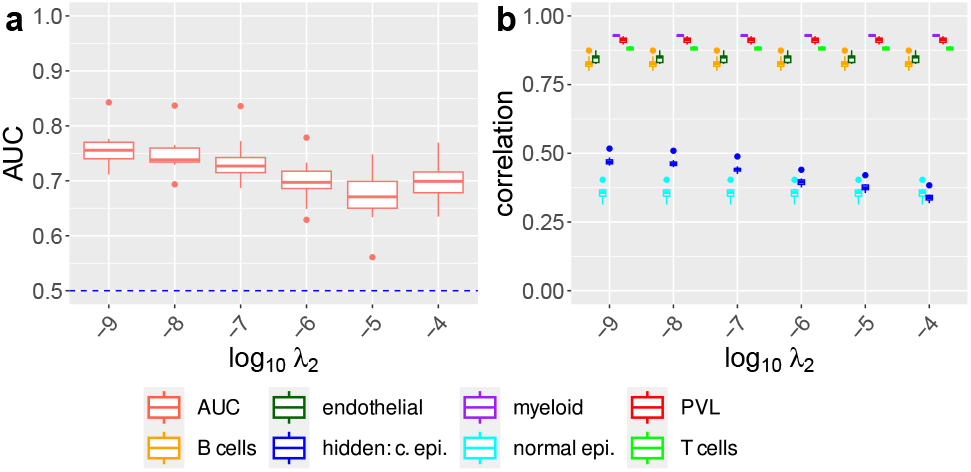
ADTD recovers cellular regulation and retains prediction performance in the breast cancer test scenario. The left Figure shows ROC AUCs for recovering cellular regulation simulated in all cell types for different regularization parameters *λ*_2_ and fixed value *λ*_1_ = 10^−1^ for ADTD in the breast cancer test study. The corresponding performance in terms of Pearson’s correlation for ADTD for estimating the known and hidden cellular contributions is shown on the right. AUCs were determined by averaging AUCs for detecting up- and down-regulated genes. Abbreviation: “hidden: c. epi.” = hidden cancer epithelial cells.

Finally, we tested how the recovery of cell-type specific regulation depends on the abundance of cell types. To study this aspect, we repeated the generation of the breast cancer test mixtures but now with (1) modified T-cells only, and (2) modified PVLs only. T cells are highly abundant, contributing at average ∼ 39.1% of cells to the mixtures, while PVLs are rare with an average contribution of 6.0%. Respective results are summarized in Suppl. Fig. S9 for T cells and in Suppl. Fig. S10 for PVLs, suggesting that ADTD infers cellular regulation irrespective of the cellular abundance.

##### Hyper-parameter dependency

while we did not utilize the test data to refine the hyper-parameters, examining their impact on the results is still insightful. We therefore evaluated the performance of ADTD for estimating the known cellular contributions, the hidden ones, and the cellular regulation for the full grid of hyper-parameters investigated previously (Fig. S2d-f). As before, considering estimates of cellular contributions, the results were stable and are compromised only if both *λ*_1_ and *λ*_2_ are very small. Moreover, estimates of cellular regulation were reasonable for a broad range of *λ*_2_ values. Thus, our results suggest a highly competitive performance of ADTD largely independent of the explicit choice of hyper-parameters, given that *λ*_2_ is in a reasonable range and that both are not simultaneously very small.

It is also worth emphasizing an analogy to other regularized regression approaches such as ordinary Lasso or ridge regression (Hoerl and Kennard, 1970; Tibshirani, 1996). Here, larger regularization parameters correspond to models which only recover the most dominant effects. As such, the user might decide whether it is important to detect the strongest regulatory effects with high confidence or the majority of regulatory effects with low confidence. Here, the former would suggest comparatively high values of *λ*_2_ and the latter comparatively low ones, corresponding to a more conservative and a more liberal choice, respectively.

### ADTD breast cancer subtype analysis using bulk transcriptomics data from TCGA

To illustrate how ADTD can complement routine data analysis, we performed an exemplary analysis considering breast cancer specimens from TCGA (see Methods). We evaluated ADTD separately for triple-negative breast cancer (TNBC), human epidermal growth factor receptor 2-positive breast cancer (HER2+), luminal A (LumA) and luminal B (LumB) breast cancer to study common and complementary cell-type specific gene regulation. Our hyper-parameters where selected as *λ*_1_ = 0.1 and 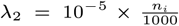, where *n*_*i*_ equals the number of samples of breast-cancer subtype *i*. The additional subtype-specific rescaling factor is necessary to make the estimated cell-type specific regulation comparable. The overall scale was motivated by the fact that values between 10^−6^ and 10^−5^ still yielded reasonable performance on the validation data, while being considerably more conservative (more regularized). We extracted the matrices Δ and denote them as Δ^(*·*)^ for breast cancer subtype (·) in the following, where the entry 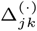 contains the respective re-scaling factor for gene *j* in cell type *k*. For instance, a ground truth 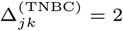 corresponds to a gene *j* being up-regulated by a factor of 2 in TNBC compared to healthy breast tissue. Possible cross-platform issues between the single-cell derived reference matrix *X* and the TCGA bulk profiles were addressed by gene-wise rescalings (see Methods).

First, we ranked the rows (genes) of Δ from highest to lowest deviations from one defined by 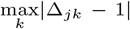. Thus, we identified the genes with strongest regulation according to ADTD for each of the four cancer subtypes across all cell types with included reference profiles. We then extracted the top ten genes for each subtype and explored the overlap (Fig. 3). We observed a remarkable overlap of 5 genes (of a total of *p* = 957 genes), which were selected in all four subtypes, namely B2M, TMSB4X, FTH1, LTB, and RPS27. We next verified if these five genes are consistently regulated among the four subtypes. B2M shows its highest regulation in myeloid cells across all subtypes, with the only exception of HER2+ breast cancers, where myeloid cells were also upregulated but slightly outcompeted by T-cells 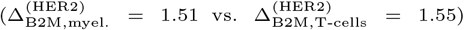. B2M encodes beta-2-microglobulin, a component of the class I major histocompatibility complex (MHC) which is present for all nucleated cells and which serves for antigen presentation. Myeloid cells comprise professional antigen presenting cells such as dendritic cells and tissue macrophages (Bassler *et al*., 2019).

**Fig. 3.**
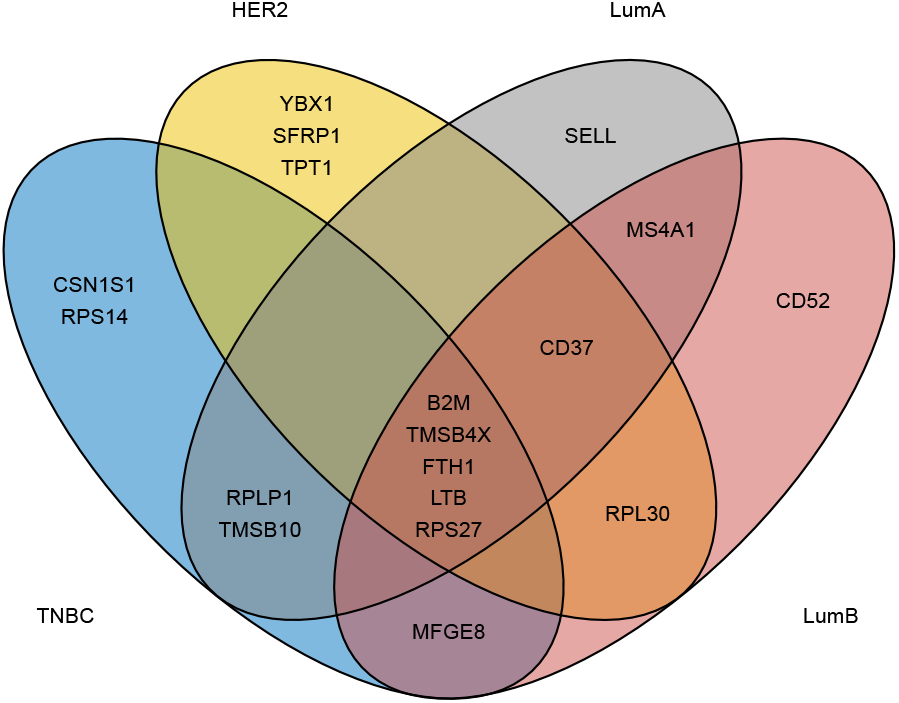
ADTD analysis of TCGA breast cancer data. The Venn diagram shows the overlap among the top ten regulated genes in negative breast cancer (TNBC), human epidermal growth factor receptor 2-positive breast cancer (HER2+), luminal A (LumA) and luminal B (LumB) breast cancer.

In line with our results, beta-2-microglobulin (*β*2-M) protein expression was significantly higher in breast cancer compared to benign breast tumors (Li *et al*., 2014). Moreover, *β*2-M protein expression was shown to be significantly different in the four breast cancer molecular subtypes (Li *et al*., 2014). ADTD adds to this observation the cell-type specific regulation of B2M, also indicating quantitative differences between TNBC, HER2+ and Luminal A/B (Figure S11a). TMSB4X encodes the protein Thymosin beta-4 and FTH1 encodes the heavy subunit of ferritin. In our analysis, both TMSB4X and FTH1 were consistently upregulated in myeloid cells across all subtypes (Figure S11e and S11f). Thymosin beta-4 was repeatedly studied in the context of breast cancer (Zhang *et al*., 2017), and in line with our findings, macrophages showed intense reactivity for Thymosin beta-4 antibodies in breast cancer, while cancer cells showed a much more variable reactivity (Larsson and Holck, 2007). Also FTH1 was repeatedly discussed in the context of breast cancer and it was shown that ferritin stimulates breast cancer cells through an iron-independent mechanism and is localized within tumor-associated macrophages (Alkhateeb *et al*., 2013). LTB encodes lymphotoxin-beta (LT-beta), also known as tumor necrosis factor C (TNF-C). LTB was consistently upregulated in B-cells (Figure S11d). B cells can produce lymphotoxin, which induces angiogenesis and thus promotes tumor growth (Yuen *et al*., 2016). Finally, RPS27 encodes ribosomal protein S27.

ADTD derived the strongest regulation in normal epithelial cells for all subtypes except HER2+. In the latter, RPS27 was slightly stronger regulated in myeloid cells 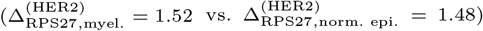, see also Figure S11c. In order to identify genes with a diverse regulation pattern derived by ADTD, we globally searched the scaling matrices Δ^(*·*)^ for largest deviations from 1. This yielded MFGE8 as top hit, observed in TNBC only. MFGE8 shows a substantial regulation of 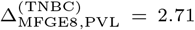 in PVLs, indicating an almost 3-fold increase while the other cell types are basically not affected (see also Figure S11b). MFGE8 encodes Milk fat globule-EGF factor 8 and its reduction was shown to inhibit triple-negative breast cancer cell viability and migration (Yang *et al*., 2019).

## Conclusion and Discussion

Multiple methods for digital tissue dissection were suggested in recent years, based on different losses such as squared residuals, *l*_1_ or the support vector regression (Altboum *et al*., 2014; Chen *et al*., 2018). Moreover, machine learning techniques such as deep learning were proposed as an alternative and promise improved performance, although adherence to strict Standard Operating Procedures (SOPs) might be necessary to achieve such a performance gain (Lin *et al*., 2022; Menden *et al*., 2020). Here, we approached computational tissue dissection from the perspective that strict SOPs are difficult (if not impossible) to guarantee and that methods should intrinsically account for between-dataset variability. As such, we proposed Adaptive Digital Tissue Deconvolution (ADTD) as an adaptive method for digital tissue dissection, which incorporates two sources of between-dataset variability. First, ADTD accounts for possibly hidden cellular background contributions, which are not represented by archetypical reference profiles. This allowed us to achieve improved out-of-sample performance as demonstrated by artificial test mixtures generated from single-cell RNA profiles of breast cancer specimens. In fact, both the proportions of background contributions and their representation as a reference profile could be estimated with confidence even for small sample sizes of *n* ∼ 100. Second, ADTD accounts for environmental effects; depending on the tissue context, cellular phenotypes might be altered (Schelker *et al*., 2017; Racle *et al*., 2017) and therefore ADTD adapts its reference matrix to the specific application. As demonstrated in comprehensive simulation studies, this allows to resolve cell-type specific regulation from bulk transcriptomics data.

It is worth emphasizing that approaching the former two issues requires appropriate strategies to deal with overfitting. Here, a common technique from high-dimensional statistics was used, namely *l*_2_ regularization (Hoerl and Kennard, 1970). Here, 𝓁_2_ regularization is not the only possible choice and was made for computational reasons, since ADTD Eq. (9) can be minimized by iteratively solving sub-problems using quadratic programming. In principle, 𝓁_1_ regularization could be an interesting alternative to enforce sparseness in the cell-regulation parameters, potentially improving the interpretability of the results. However, to further explore this option would require appropriate implementation strategies. It is also important to point out that tuning the strength of the two *l*_2_-regularization terms is achieved by adjusting two hyper-parameters *λ*_1_ and *λ*_2_. Typically, hyper-parameters are selected by lowest generalization error. This strategy, however, fails for cell-type deconvolution; we are not interested in a low generalization error for the prediction of the bulk profiles *Y*, but in low errors of the underlying parameters *C, c*, and *x*. Therefore, we decided to perform hyper-parameter selection via a controlled validation study where the ground truth is available. Surprisingly, results were rather insensitive to the explicit choice of *λ*_1_ and *λ*_2_, under the assumption that both are not too small simultaneously. This means that *a priori* parameter choices might be appropriate for most applications. Alternatively, one might select hyper-paramters as follows: (A) start out with comparatively high regularization parameters. This typically yields reliable results for estimating cellular proportions, while the cell-type specific gene regulation cannot be resolved. Then, (B) systematically lower *λ*_2_ to resolve gene regulation. While doing so, respective estimates for cellular proportions should be controlled for deviations from the previous results. In our studies, the improvements from further tuning *λ*_1_ were negligible and thus it could remain fixed.

It is noteworthy that recent deep-learning based approaches can address more complex non-linear relationships. An example is Scaden, which, although it performed substantially better than CIBERSORTx and EPIC, could not outperform ADTD. The underlying reason could be that bulk profiles are inherently linear, as they correspond to a linear combination of individual cells’ contributions. Thus, a bias towards linearity could be beneficial in this context. Finally, cell-type deconvolution is designed to disentangle molecularly diverse cells and performance is typically compromised in scenarios where many cell types are disentangled. This is particularly the case if related cell types are considered. We demonstrated that the adapted deconvolution weights *g* in DTD, which are also included in ADTD, improve on this issues (Görtler *et al*., 2020). However, still it is insufficiently addressed given the fact that molecularly similar cells can fulfill highly diverse tasks.

In summary, ADTD represents a substantial step in the development of digital tissue dissection methods, as it is to the best of our knowledge the first method which tackles two big intrinsic model uncertainties simultaneously, namely hidden background contributions and cellular environmental factors. These two aspects might not just be relevant for the analysis of bulk transcriptomics data, but also for new data data sources, such as spatial transcriptomics data. Also there, accounting for domain transfer and hidden variables, such as cellular background contributions, might be key for reliable inference.

