## Supplement for "Adaptive Digital Tissue Deconvolution"

### Supplementary Material: Adaptive Digital Tissue Deconvolution

#### 1 Algorithmic details

##### 1.1 Estimate hidden cell proportions

The hidden cell proportions $c = (c_1, c_2, \ldots, c_n) \in \mathbb{R}_+^{1\times n}$ of ADTD can be estimated analogously to Eq. (7) of the main manuscript as follows:

$$
\begin{aligned}
Y_{\cdot,i} &= \Delta X C_{\cdot,i} + x c_i + \epsilon_{\cdot,i} \\
&\Rightarrow \quad 1 = \sum_{j=1}^{p} Y_{ji} = \sum_{j=1}^{p}\sum_{k=1}^{q} \Delta_{jk} X_{jk} C_{ki} + \sum_{j=1}^{p} x_j c_i + \epsilon_i \\
&\Rightarrow \quad c_i = 1 - \sum_{j=1}^{p}\sum_{k=1}^{q} \Delta_{jk} X_{jk} C_{ki} \\
&\Rightarrow \quad c = \max\left((0, \cdots, 0), J_{1,n} - J_{1,p}(\Delta \circ X) C\right), \qquad (1)
\end{aligned}
$$

where we used $\sum_{j=1}^{p} x_j = 1$ and assumed that the $\epsilon_i$ are small.

##### 1.2 Optimization with respect to $C$

Next, we derive an estimate for $C$. Consider the ADTD loss function $L_{\text{ADTD}}(C, x, \Delta)$ for given $x$ and $\Delta$:

$$
\begin{aligned}
F(C) &= L_{\text{ADTD}}(C, x, \Delta) \\
&= ||G\left(Y - (\Delta \circ X)C - x\left(J_{1,n} - J_{1,p}(\Delta \circ X)C\right)\right)||_F^2 + \lambda_1 ||C_0 - C||_F^2 + const. \\
&= ||GY - GxJ_{1,n} - G\left(\Delta \circ X - xJ_{1,p}(\Delta \circ X)\right)C||_F^2 + \lambda_1 ||C_0 - C||_F^2 + const. \\
&= \left|\left| \begin{pmatrix} GY - GxJ_{1,n} \\ \sqrt{\lambda_1} C_0 \end{pmatrix} - \begin{pmatrix} G\left(\Delta \circ X - xJ_{1,p}(\Delta \circ X)\right) \\ \sqrt{\lambda_1} I_q \end{pmatrix} C \right|\right|_F^2 + const. \qquad (2)
\end{aligned}
$$

The latter expression allows us to reformulate the estimate of $C$ as a quadratic programming problem. Let $\mathbf{c} = C_{\cdot i}$ and $y = Y_{\cdot i}$, then estimating $C$ can be achieved by minimizing

$$\frac{1}{2}\mathbf{c}^T P \mathbf{c} + q^T \mathbf{c} \qquad (3)$$

subject to

$$J_{1,p}(\Delta \circ X)\mathbf{c} \leq (1, \ldots, 1)$$

and

$$-\mathbf{c} \preceq 0\,.$$

with respect to $\mathbf{c}$, where

$$P = 2\begin{pmatrix} G\left(\Delta \circ X - xJ_{1,p}(\Delta \circ X)\right) \\ \sqrt{\lambda_1}I_q \end{pmatrix}^T \begin{pmatrix} G\left(\Delta \circ X - xJ_{1,p}(\Delta \circ X)\right) \\ \sqrt{\lambda_1}I_q \end{pmatrix}\,,$$

$$Q^T = -2\begin{pmatrix} GY - GxJ_{1,n} \\ \sqrt{\lambda_1}C_0 \end{pmatrix}^T \begin{pmatrix} G\left(\Delta \circ X - xJ_{1,p}(\Delta \circ X)\right) \\ \sqrt{\lambda_1}I_q \end{pmatrix}$$

and

$$q = Q_{\cdot i}\,.$$

This procedure is performed for all columns $C_{\cdot i}$.

#### 1.3 Optimization with respect to $x$

Finally, we derive an estimate for $x$, where we use the abbreviation $Z = Y - \Delta XC$:

$$\begin{aligned}
L_{\text{ADTD}}(x) &= ||GZ - Gxc||_F^2 \\
&= \operatorname{Tr}\left[(Z - xc)^T\Gamma(Z - xc)\right] \\
&= \operatorname{Tr}\left[c^T x^T \Gamma xc\right] - 2\operatorname{Tr}\left[Z^T\Gamma xc\right] \\
&= \operatorname{Tr}\left[cc^T x^T \Gamma x\right] - 2\operatorname{Tr}\left[cZ^T\Gamma x\right] \\
&= (cc^T)(x^T\Gamma x) - 2cZ^T\Gamma x \\
&= \frac{1}{2}x^T P' x + q'^T x
\end{aligned}$$

subject to $x \succeq 0$ and $\sum_{j=1}^p x_j = 1$, with $P' = 2(cc^T)\Gamma$ and $q'^T = -2cZ^T\Gamma$, where one should note that $cc^T$ and $x^T\Gamma x$ are scalars. Thus, also this optimization problem reduces to quadratic programming.

#### 1.4 Optimization with respect to $\Delta$

In the following, we derive a procedure to minimize $L_{\text{ADTD}}(C, x, \Delta)$ with respect to $\Delta_{j,\cdot}$, while $C$, $x$ and $\Delta_{k,\cdot}$ with $k \neq j$ are kept fixed. Let, $\delta_k = (0, \ldots, 0, 1, 0 \ldots, 0)^T$, where the 1 is at the $k$th position and let $\delta_{\neq k} = (1, \ldots, 1)^T -$

$\delta_k$. Consider $F(\Delta_{j,\cdot}) = L_{\text{ADTD}}(C, x, \Delta)$. Then $F(\Delta_{j,\cdot})$ becomes

$$\begin{aligned}
& ||G\left(Y - xJ_{1,n} - (\Delta \circ X)C + xJ_{1,p}(\Delta \circ X)C\right)||_F^2 + \lambda_2 ||\Delta - J_{p,q}||_F^2 \\
= \; & ||G_{jj}\Big(Y_{j,\cdot} - x_j J_{1,n} + x_j \delta_{\neq j}^T (\Delta \circ X)C - (\Delta \circ X)_{j\cdot}C + x_j \delta_j^T (\Delta \circ X)C\Big)||_F^2 \\
& + \sum_{k \neq j} ||G_{kk}\Big(Y_{k,\cdot} - x_k J_{1,n} - (\Delta \circ X)_{k,\cdot}C + x_k \delta_{\neq j}(\Delta \circ X)C + x_k \delta_j (\Delta \circ X)C\Big)||_F^2 \\
& + \lambda_2 ||\Delta_{j\cdot} - J_{1,q}||_F^2 + const. \\
= \; & ||G_{jj}\left(A_{j,\cdot} - (1 - x_j)(\Delta \circ X)_{j,\cdot}C\right)||_F^2 + \sum_{k \neq j} ||G_{kk}\Big(B_{k,\cdot} + x_k \Delta_{j,\cdot}(\Delta \circ X)_{k,\cdot}C\Big)||_F^2 \\
& + \lambda_2 ||\Delta_{j,\cdot} - J_{1,q}||_F^2 + const. \\
= \; & ||G_{jj}\left(A_{j,\cdot} - (1 - x_j)\Delta_{j,\cdot}C_{X_{j,\cdot}}\right)||_F^2 + \sum_{k \neq j} ||G_{kk}\Big(B_{k,\cdot} + x_k \Delta_{j,\cdot}C_{X_{j,\cdot}}\Big)||_F^2 \\
& + \lambda_2 ||\Delta_{j,\cdot} - J_{1,q}||_F^2 + const.\,,
\end{aligned}$$

with

$$\begin{aligned}
A_{j,\cdot} &= Y_{j,\cdot} - x_j J_{1,n} + x_j \delta_{\neq j}^T (\Delta \circ X)C\,, \\
B_{k,\cdot} &= Y_{k,\cdot} - x_k J_{1,n} - (\Delta \circ X)_{k,\cdot}C + x_k \delta_{\neq j}(\Delta \circ X)C\,,
\end{aligned} \tag{4}$$

where we used the abbreviation $C_{X_{j,\cdot}} = (X_{j,\cdot}^T \circ C_{\cdot,1}, \dots, X_{j,\cdot}^T \circ C_{\cdot,n})$ and summarized all terms independent of $\Delta_{j,\cdot}$ as *const.*. To minimize the former equation using quadratic programming, we rewrite it as

$$\begin{aligned}
F(\Delta_{j,\cdot}) = \; & G_{jj}^2 \left(A_{j,\cdot}A_{j,\cdot}^T - 2(1 - x_j)A_{j,\cdot}C_{X_{j,\cdot}}^T \Delta_{j,\cdot}^T + (1 - x_j)^2 \Delta_{j,\cdot}C_{X_{j,\cdot}}C_{X_{j,\cdot}}^T \Delta_{j,\cdot}^T\right) \\
& + \sum_{k \neq j} G_{kk}^2 \left(B_{k,\cdot}B_{k,\cdot}^T + 2x_k B_{k,\cdot}C_{X_{j,\cdot}}^T \Delta_{j,\cdot}^T + x_k^2 \Delta_{j,\cdot}C_{X_{j,\cdot}}C_{X_{j,\cdot}}^T \Delta_{j,\cdot}^T\right) \\
& + \lambda_2 \left(J_{1,q}J_{1,q}^T - 2J_{1,q}\Delta_{j,\cdot}^T + \Delta_{j,\cdot}\Delta_{j,\cdot}^T\right)\,.
\end{aligned} \tag{5}$$

Let $\mathbf{b} = \Delta_{j,\cdot}^T$,

$$P = 2\left(G_{jj}^2(1 - x_j)^2 + \sum_{k \neq j} G_{kk}^2 x_k^2\right) C_{X_{j,\cdot}}C_{X_{j,\cdot}}^T + 2\lambda_2 I_q$$

and

$$q^T = -2\left(G_{jj}^2(1 - x_j)A_{j,\cdot} - \sum_{k \neq j} G_{kk}^2 x_k B_{k,\cdot}\right) C_{X_{j,\cdot}}^T - 2\lambda_2 J_{1,q}\,,$$

to formulate a typical quadratic programming problem:

$$F(\mathbf{b}) = \frac{1}{2}\mathbf{b}^T P \mathbf{b} + q^T \mathbf{b} \tag{6}$$

subject to constraints

$$\mathbf{b} \succeq (0, \ldots, 0)^T, \quad \text{and} \quad J_{1,n} - \delta_{\neq k}^T (\Delta \circ X) C \succeq \mathbf{b}^T C_{X_{k,\cdot}}\,. \tag{7}$$

The latter constraint can be derived from ADTD constraint $J_{1,q}(\Delta \circ X)C \preceq J_{1,n}$.

### 2 Supplementary Figures and Tables

Table S1: **Performance of ADTD, EPIC, CIBERSORTx and Scaden on training data.** Observed Pearson's correlations obtained by comparing the estimated cellular proportions with the ground truth for artificial cellular training mixtures generated from single-cell data of healthy breast tissue specimens (see Methods). The errors correspond to ±1 standard deviation obtained over 10 simulation runs. For ADTD the parameters $\lambda_1 = 10^{-1}$ and $\lambda_2 = 10^{-8}$ were used (see hyper-parameter selection in validation performance section). Abbreviations: Endo. = endothelial cells; Myel. = myeloid cells; Epith. = epithelial cells; PVL = perivascular-like cells

| | ADTD | $EPIC_1$ | $EPIC_2$ |
|---|---|---|---|
| B-cells | 0.791 ± 0.020 | 0.112 ± 0.031 | 0.648 ± 0.014 |
| Endo. | **0.941 ± 0.006** | **0.863 ± 0.005** | 0.672 ± 0.015 |
| Myel. | **0.943 ± 0.005** | **0.854 ± 0.007** | 0.687 ± 0.015 |
| Epith. | **0.888 ± 0.008** | 0.647 ± 0.014 | - |
| PVL | **0.894 ± 0.011** | 0.573 ± 0.019 | - |
| T-cells | **0.942 ± 0.003** | **0.813 ± 0.009** | 0.293 ± 0.02 |
| mean | **0.900 ± 0.003** | 0.644 ± 0.007 | 0.577 ± 0.008 |

| | CIBERSORTx | Scaden |
|---|---|---|
| B-cells | 0.179 ± 0.041 | 0.486 ± 0.025 |
| Endo. | **0.825 ± 0.007** | 0.782 ± 0.012 |
| Myel. | **0.87 ± 0.006** | **0.812 ± 0.015** |
| Epith. | **0.806 ± 0.008** | 0.783 ± 0.008 |
| PVL | 0.707 ± 0.012 | 0.756 ± 0.01 |
| T-cells | **0.826 ± 0.009** | 0.715 ± 0.016 |
| mean | 0.702 ± 0.006 | 0.722 ± 0.006 |

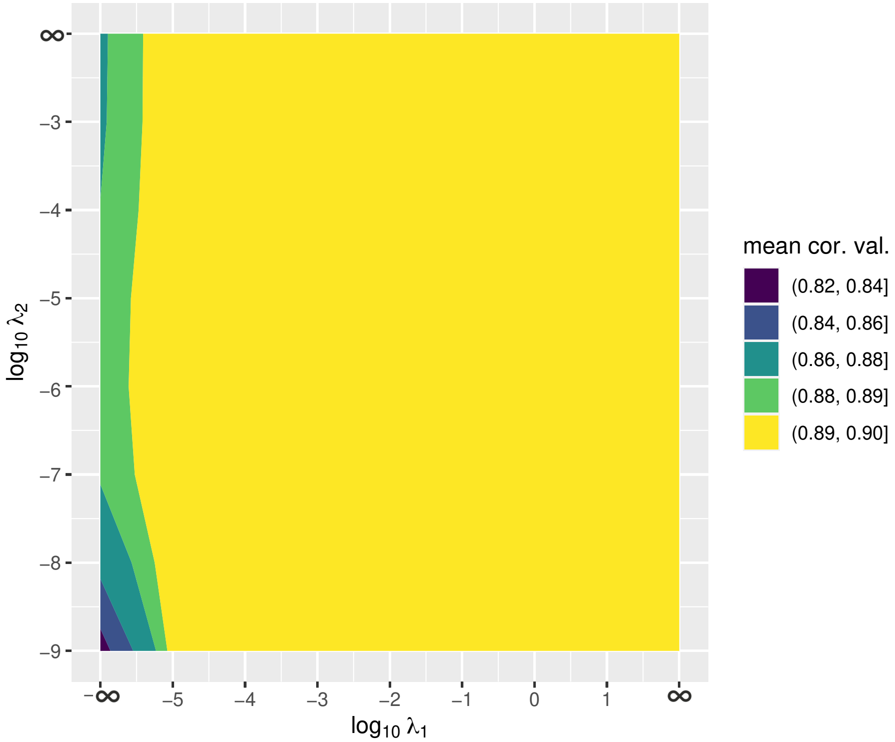


Figure S1: **ADTD performance for different hyper parameters on the training data.** A comprehensive parameter grid consisting of all combinations of $\lambda_1 \in \{0, 10^{-5}, 10^{-4}, \ldots, 1, 10, \infty\}$ with $\lambda_2 \in \{10^{-9}, 10^{-8}, \ldots, 10, \infty\}$ was evaluated. Performance was assessed by (1) calculating Pearson's correlation between ground truth and predictions for each of the included cell types, and (2) by subsequently averaging them.

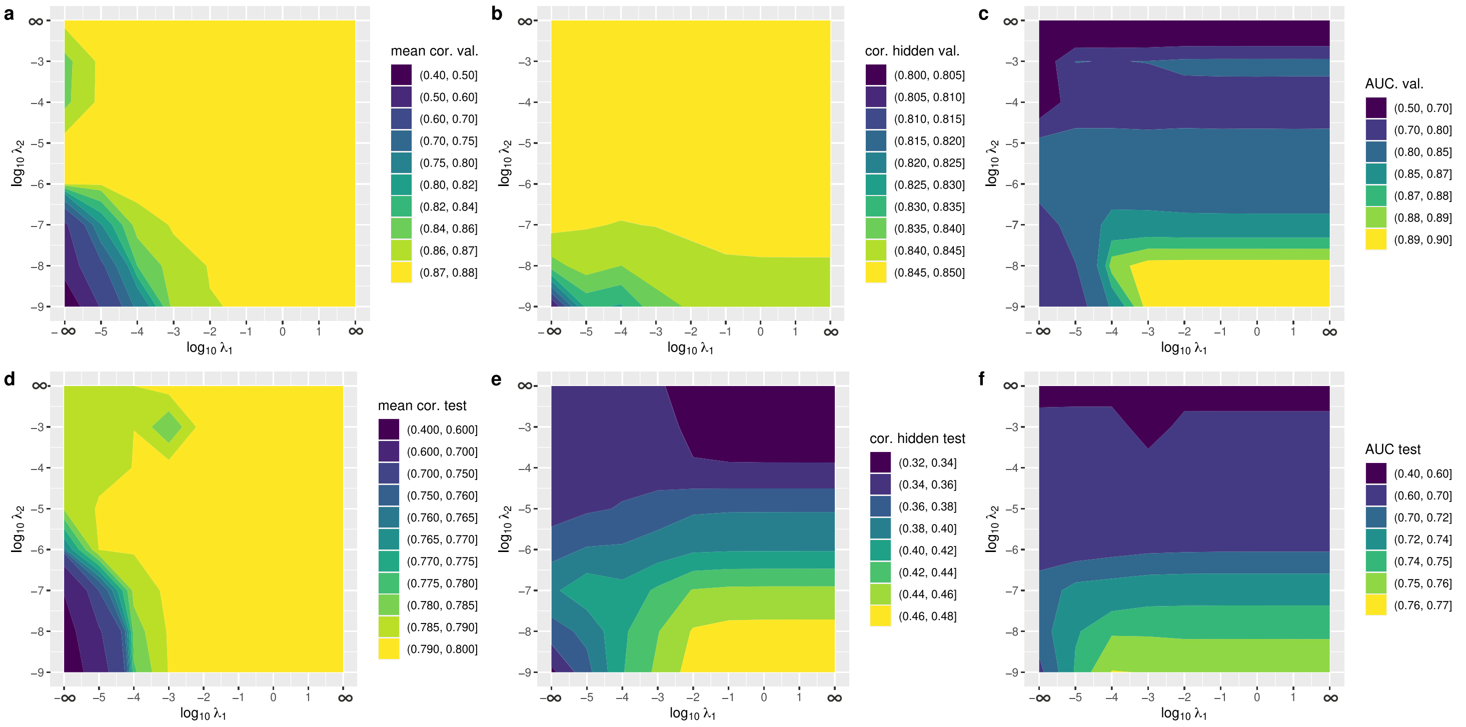


Figure S2: **ADTD performance for different hyper parameters on the validation and test data data.** A comprehensive parameter grid consisting of all combinations of $\lambda_1 \in \{0, 10^{-5}, 10^{-4}, \dots, 1, 10, \infty\}$ with $\lambda_2 \in \{10^{-9}, 10^{-8}, \dots, 10, \infty\}$ was evaluated. Figure a to c correspond to the validation data and d to e to the test data. Figure a and d show the average performance in predicting the known cellular contributions, and b and e for the hidden contributions. Fig. c and f give the corresponding areas under the ROC curves for detecting cell-type specific gene regulation.

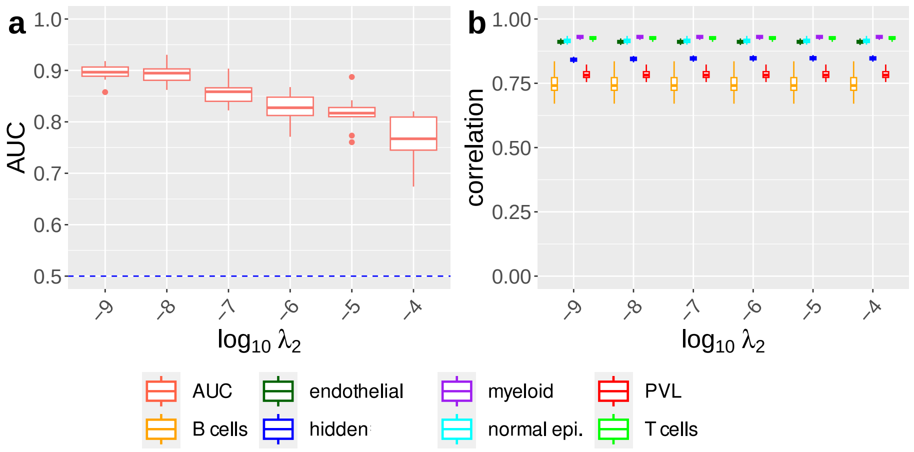


Figure S3: **ADTD performance for recovering cell-type specific gene regulation on the validation data.** The left figure shows areas under the ROC curve for recovering cellular regulation for different regularization parameters $\lambda_2$, where $\lambda_1 = 10^{-1}$ was kept fixed. The corresponding performance in terms of Pearson's correlation for ADTD for estimating the known and hidden cellular contributions is shown on the right. Abbreviation: "hidden" = hidden cellular contributions.

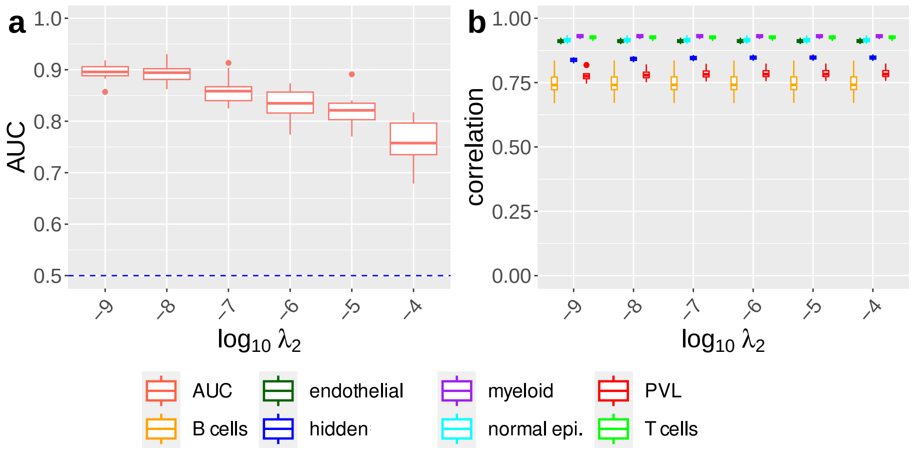


Figure S4: **ADTD performance for recovering cell-type specific gene regulation on the validation data.** The left figure shows areas under the ROC curve for recovering cellular regulation for different regularization parameters $\lambda_2$, where $\lambda_1 = 10^{-3}$ was kept fixed. The corresponding performance in terms of Pearson's correlation for ADTD for estimating the known and hidden cellular contributions is shown on the right. Abbreviation: "hidden" = hidden cellular contributions.

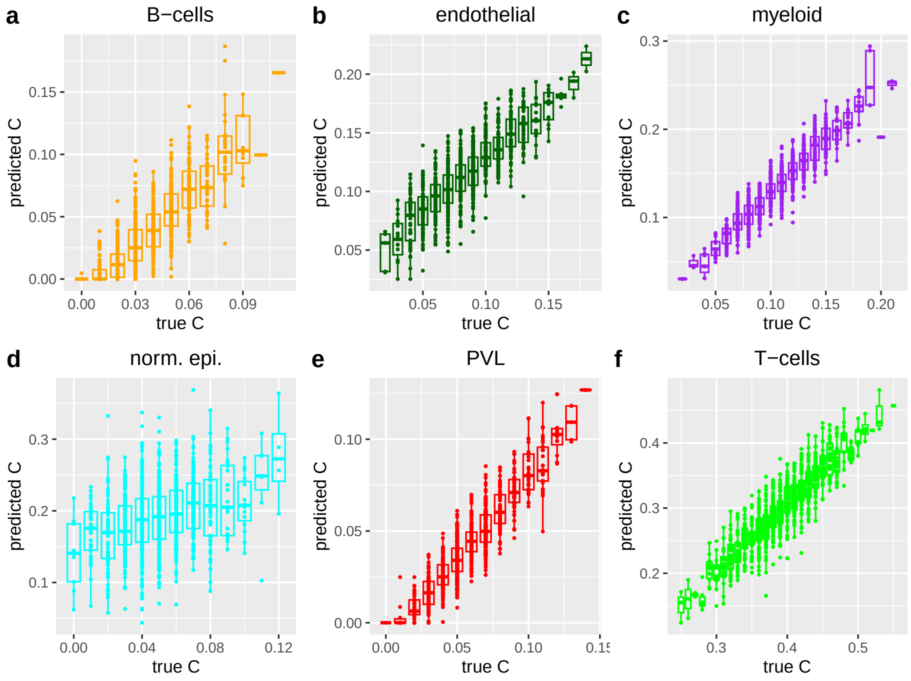


Figure S5: **Visualization of ADTD results on test data.** Predicted versus true cellular composition for the cell types captured in the reference matrix $X$ for one simulation run of the testing scenario ($n = 1000$).

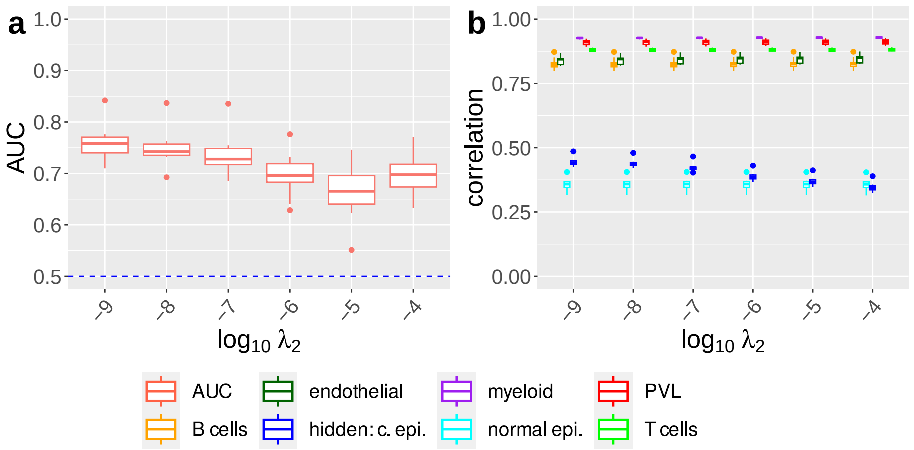


Figure S6: **ADTD performance for recovering cell-type specific gene regulation on the breast cancer test data.** The left figure shows areas under the ROC curve for recovering cellular regulation for different regularization parameters $\lambda_2$, where $\lambda_1 = 10^{-3}$ was kept fixed. The corresponding performance in terms of Pearson's correlation for ADTD for estimating the known and hidden cellular contributions is shown on the right. Abbreviation: "hidden: c. epi." = hidden cancer epithelial cells.

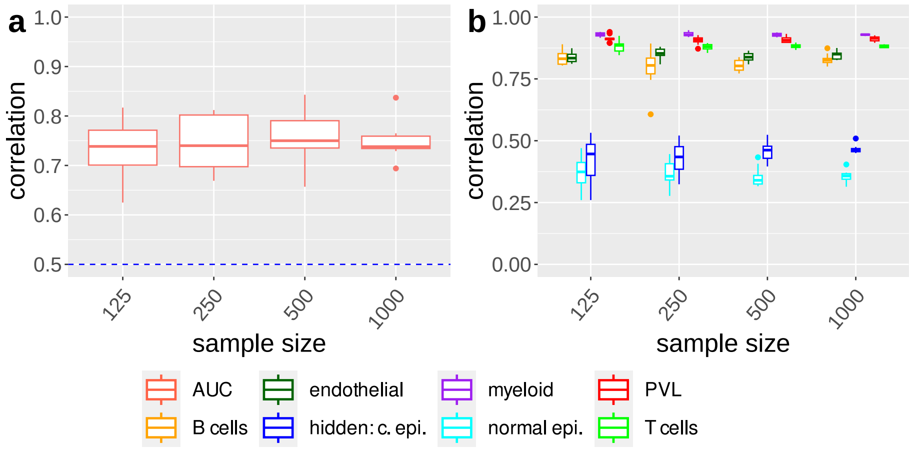


Figure S7: **ADTD performance versus sample size in the breast cancer test data.** Figure a shows areas under the ROC curve for recovering cellular regulation for different sample sizes $n = 125, 250, 500, 1000$ for ADTD ($\lambda_1 = 10^{-1}$, $\lambda_2 = 10^{-8}$). The corresponding performance in terms of Pearson's correlation for estimating the known and hidden cellular contributions is shown on the right. Abbreviation: "hidden: c. epi." = hidden cancer epithelial cells.

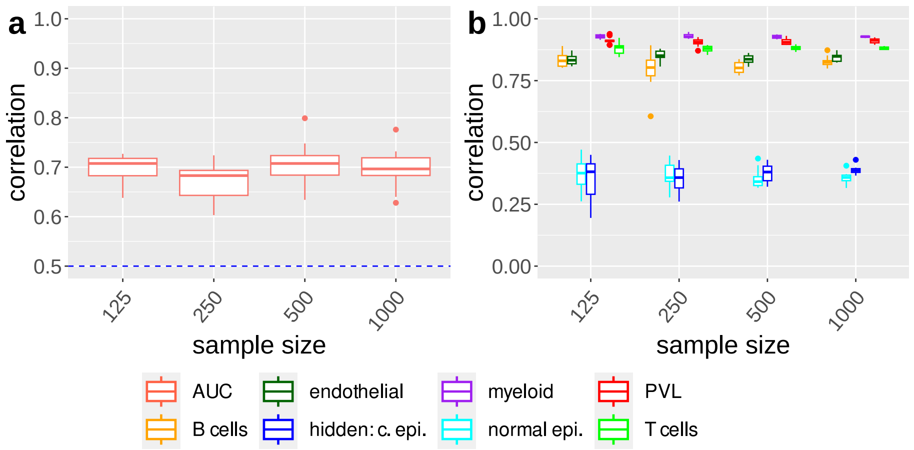


Figure S8: **ADTD performance versus sample size in the breast cancer test data.** Figure a shows areas under the ROC curve for recovering cellular regulation for different sample sizes $n = 125, 250, 500, 1000$ for ADTD ($\lambda_1 = 10^{-3}$, $\lambda_2 = 10^{-6}$). The corresponding performance in terms of Pearson's correlation for estimating the known and hidden cellular contributions is shown on the right. Abbreviation: "hidden: c. epi." = hidden cancer epithelial cells.

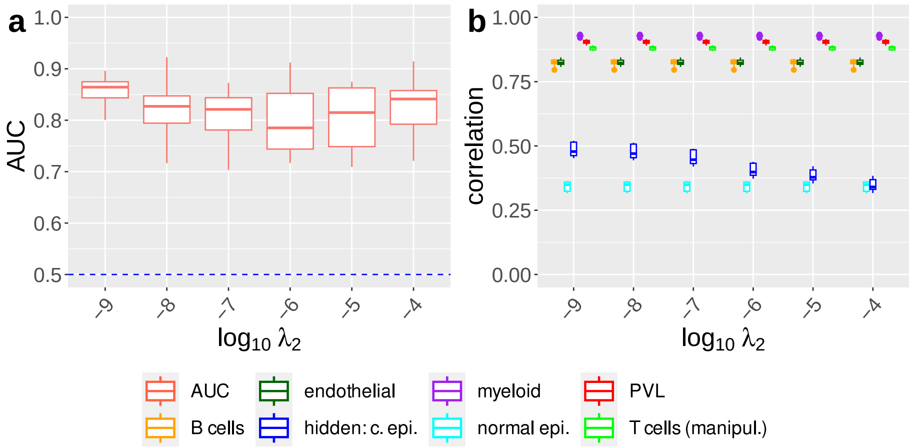


Figure S9: **ADTD performance for recovering cell-type specific gene regulation in T-cells on the breast cancer test data.** The left figure shows areas under the ROC curve for recovering cellular regulation in T cells for different regularization parameters $\lambda_2$, where $\lambda_1 = 10^{-1}$ was kept fixed. The corresponding performance in terms of Pearson's correlation for ADTD for estimating the known and hidden cellular contributions is shown on the right. Abbreviation: "hidden: c. epi." = hidden cancer epithelial cells.

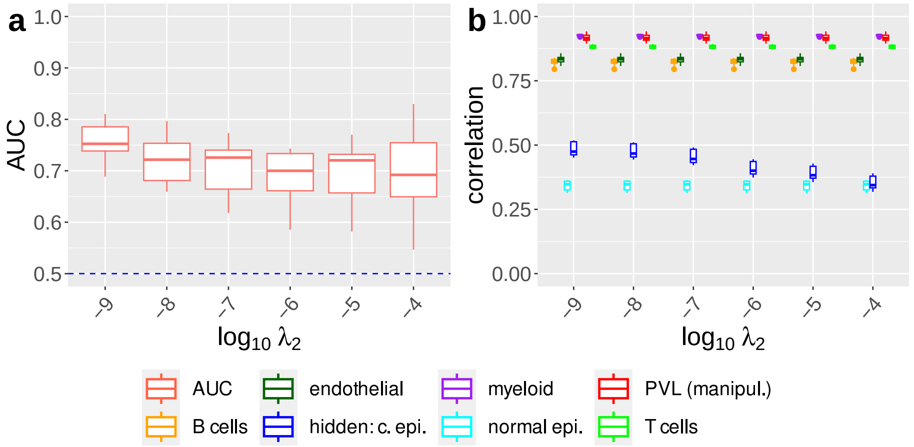


Figure S10: **ADTD performance for recovering cell-type specific gene regulation in PVL cells on the breast cancer test data.** The left figure shows areas under the ROC curve for recovering cellular regulation in PVL cells for different regularization parameters $\lambda_2$, where $\lambda_1 = 10^{-1}$ was kept fixed. The corresponding performance in terms of Pearson's correlation for ADTD for estimating the known and hidden cellular contributions is shown on the right. Abbreviation: "hidden: c. epi." = hidden cancer epithelial cells

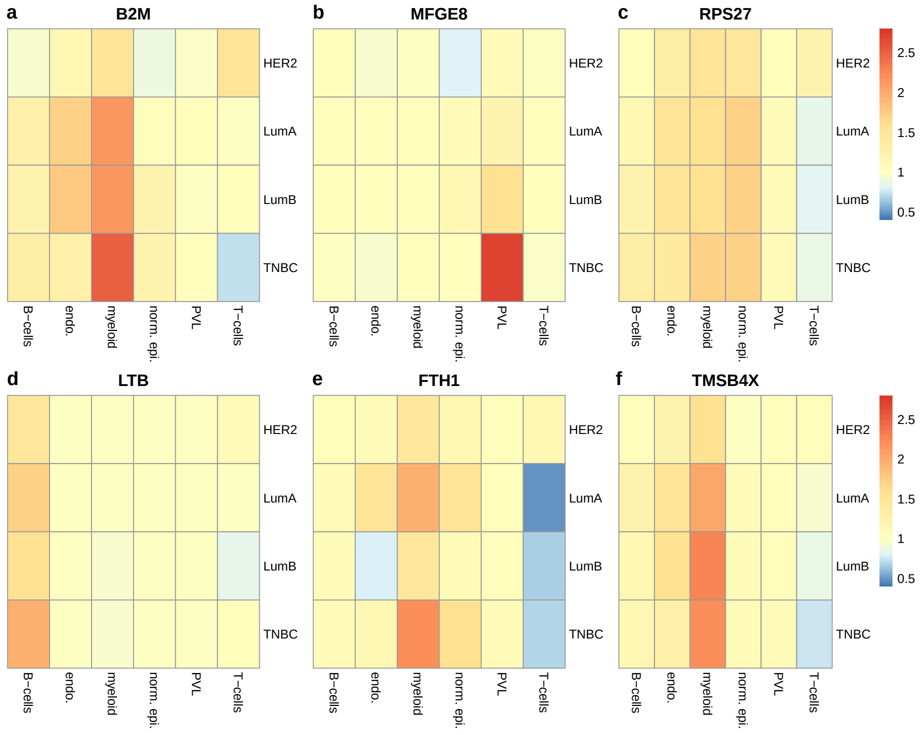


Figure S11: Δ **matrices representing cell-type specific gene regulation for six different genes discussed in the main text.** Upregulation corresponds to red and downregulation to blue colors.
